# COPO: a metadata platform for brokering FAIR data in the life sciences

**DOI:** 10.1101/782771

**Authors:** Anthony Etuk, Felix Shaw, Alejandra Gonzalez-Beltran, David Johnson, Marie-Angélique Laporte, Philippe Rocca-Serra, Elizabeth Arnaud, Medha Devare, Paul J Kersey, Susanna-Assunta Sansone, Robert P Davey

**Affiliations:** Earlham Institute, Norwich Research Park, UK; Oxford e-Research Centre, Department of Engineering Science, University of Oxford, UK; Science and Technology Facilities Council, Harwell, UK; Department of Informatics and Media, Uppsala University, Sweden; Bioversity International, France; International Food Policy Research Institute, USA; EMBL-The European Bioinformatics Institute, UK; Royal Botanic Gardens, Kew, UK

**Author notes:** Joint corresponding authors.

## Abstract

Scientific innovation is increasingly reliant on data and computational resources. Much of today’s life science research involves generating, processing, and reusing heterogeneous datasets that are growing exponentially in size. Demand for technical experts (data scientists and bioinformaticians) to process these data is at an all-time high, but these are not typically trained in good data management practices. That said, we have come a long way in the last decade, with funders, publishers, and researchers themselves making the case for open, interoperable data as a key component of an open science philosophy. In response, recognition of the FAIR Principles (that data should be Findable, Accessible, Interoperable and Reusable) has become commonplace. However, both technical and cultural challenges for the implementation of these principles still exist when storing, managing, analysing and disseminating both legacy and new data.

COPO is a computational system that attempts to address some of these challenges by enabling scientists to describe their research objects (raw or processed data, publications, samples, images, etc.) using community-sanctioned metadata sets and vocabularies, and then use public or institutional repositories to share it with the wider scientific community. COPO encourages data generators to adhere to appropriate metadata standards when publishing research objects, using semantic terms to add meaning to them and specify relationships between them. This allows data consumers, be they people or machines, to find, aggregate, and analyse data which would otherwise be private or invisible. Building upon existing standards to push the state of the art in scientific data dissemination whilst minimising the burden of data publication and sharing.

**Availability:** COPO is entirely open source and freely available on GitHub at https://github.com/collaborative-open-plant-omics. A public instance of the platform for use by the community, as well as more information, can be found at <u>copo-project.org</u>.

## Introduction

### The importance of metadata

Metadata, defined as “data about data”, provides contextual information about data and is one of the central concepts required to allow the structuring of datasets in a way that allows automated search, query and retrieval. In the life sciences, this process is called *biocuration^1^*.

Naively, humans tend to catalogue information within computer systems (and elsewhere) in ways that require subsequent interpretation. For example, naming a file `“20190114_final-with-corrections.txt”` reminds the author of certain properties of the data inside the file with very little overhead. Documents ending in the suffix “.txt”, are generally written in plain text, but this structure is explicitly given, and must be assumed by the person wanting to reuse this data. Now consider a file called `“dogs.csv”` in a folder called `“behaviour”` in a computer file system. The file has the “*csv*” extension, which stands for “comma separated values” and implies a defined tabular data structure. The file and directory names suggest that the contents relate to some kind of dog behaviour. The contents of these files cannot be fully understood until they are opened and interpreted. The representation of the information conveyed in both these examples has many ambiguities that can only be resolved by *a priori* knowledge. Without explicit boundaries and rules that describe this kind of data, misinterpretation may be commonplace and if the data is not as expected, it is impossible to know if this is because the data is corrupted or the assumptions of the finder are misplaced. Such descriptive boundaries are called *context*. Providing structured metadata makes the context for a given piece of data explicit.

Consider a table in a database called “*dog_behaviour_observations*” which keeps behavioural observations in tabular (row and column) form. Each row of the table represents a single observation, and each column represents a variable about that observation, such as “*behaviour*_*observation_id*”, “*behaviour_type*”, “*breed*”, “*date_observed*”, “*video_link*”, “*duration*”. Entries within this database now have a structure to the data, and researchers generating observation data have a (relatively) strict format to adhere to when providing data to the resource. However, this methodology still has issues. The names of the variables (column headers in the database) may not be standardised or self-describing, e.g. the variable “*duration*” might relate to an observation in a number of ways, but without further documentation, it is impossible to know this for sure. “*Duration*” may also be measured in different ways with different units, and the processes of measuring the duration of an observation may not have been recorded or standardised. Furthermore, without a way of validating the input to this database, errors may find their way in. Whilst databases do have a minimal form of validation in the form of field data types (e.g. “*behaviour_observation_id*” might be required to be a non-negative whole number, “*video_link*” a string of fewer than 255 characters, etc,), these constraints provide almost no semantic insight into the *meaning* of a field. This type of validation also does not take broader context into account, adding sparse metadata to a dataset. By using well-defined vocabulary terms for field names and values, we can easily provide information such as:

- What was the breed of the dog, based on an agreed set of dog breed names?
- How were the researchers measuring the impact of a given behaviour on other dogs, based on an agreed measurement technique?
- What was the weather like on the day of the test, based on standard meteorological descriptions?
- Are the individuals observed related, based on a common vocabulary of familial relationships?

The ability to ask and answer questions such as these may well affect the ability of another researcher to find and reuse this dataset. If we wanted to get all behavioural observations collected on a rainy day, between siblings that are all black Labradors, we couldn’t do this with our current database structure. Therefore, effective reuse of data requires that metadata be structured and composed of defined terms according to common agreed standards. Employing such standards enhances the value (and ultimate impact) of the data produced ^1^.

### The role of ontologies for interoperability

Common vocabularies, standards, and dictionaries of agreed terms, or “facts”, can be represented as ontologies, i.e. formal descriptions of a domain of knowledge ^2^. Ontologies define *axioms*, i.e. terms and the relationships between them, that can be used to construct more complex statements of knowledge. For example, we are interested in finding information on crop trials involving Chinese Spring wheat in sub-Saharan Africa. Our first problem is one of vocabulary. There is not a single shared name for this geographical region: we could search for “sub-saharan Africa”, “Africa (sub-saharan)”, or one of many other permutations. The name of the data field containing this information could also be varied e.g. “Location”, “Region”, or “Country”. If trial data is not standardised across experiments, aggregating searches across multiple datasets quickly becomes a time-consuming manual task, if not impossible. However, by labelling with unambiguous ontology terms, we can specify exactly what we mean, thus increasing the usefulness of our data to other researchers.

Figure 1 shows an example of multiple datasets being aggregated in such a way. Three columns from three different spreadsheets are shown, where the field names “**Location**”, “**region**” and “**LOC**” could be resolved by a curator to the same ontology term “**LOCATION**”, telling us that they are labelling the same entity, i.e. a geographical area. The values “**Great Rift Valley**” and “**GRV**” also resolve to the same term, meaning that a common vocabulary is used removing ambiguity. Similarly, incomplete and erroneous data can also be corrected to a degree. For instance, the value “**centra;**” would likely be assumed to be both incomplete and misspelt when assessed by a curator. However, resolving it to the term “**CENTRAL AFRICA**” overcomes these inconsistencies. To summarise, data is often messy and needs active curation to resolve issues, either through manual editing or through the use of specific software. Tools such as OpenRefine ^3^ are designed to facet human-readable terms and quickly highlight discrepancies.

**Figure 1.**
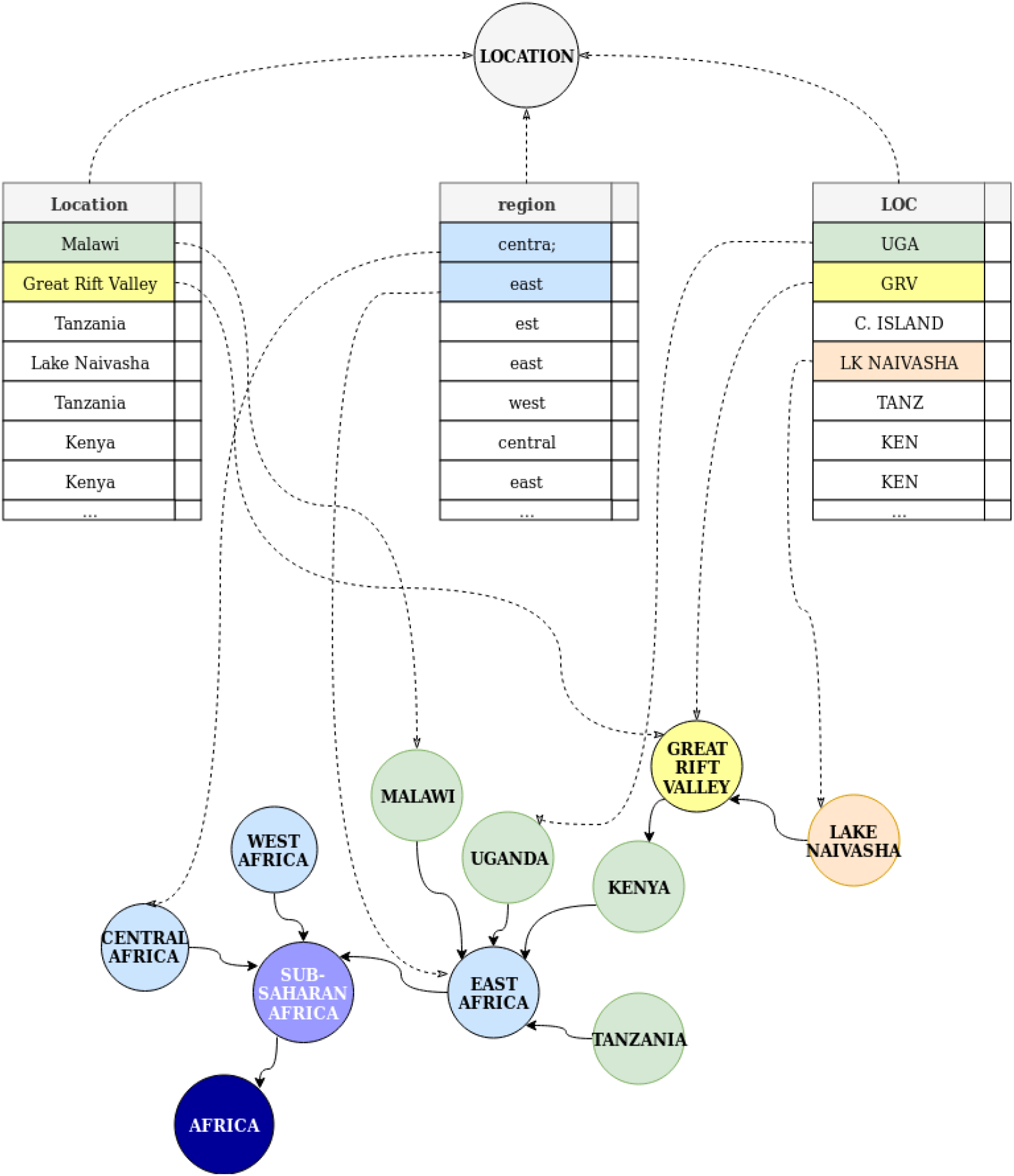
Different spreadsheet column headings can be mapped to unifying standard terms, and observations in rows can be mapped to a graph of hierarchical axioms that define a specific geographical location.

Ontology terms are typically arranged into a Directed Acyclic Graph (DAG) structure based on a model of a domain, where the nodes are the entities and the edges are the properties or relations between the entities. Part of the model may include a hierarchy, which allows us to infer context even when it is not explicitly provided. To continue the example, although none of the data in Figure 1 are labelled with the term “**Sub-Saharan Africa**”, through the contextual hierarchy of the ontology (the graph defines “**Sub-Saharan Africa**” as composed of three more specific terms “**Central Africa**”, “**West Africa**”, and **“East Africa**”) a human or a machine is able to easily infer that all these data are part of “**Sub-Saharan Africa**”. Likewise, countries in East Africa (e.g. Malawi, Uganda, Kenya, Tanzania from the example below) are also inferred to be part of Sub-Saharan Africa, providing a broader context without the need for its explicit specification, and the added effort that would require.

### Standards for data sharing

The use of ontology terms to mark up data as in the examples above means that information becomes less ambiguous, and less ambiguity makes data more Findable, Accessible, Interoperable, and Reusable (FAIR) ^4^. The FAIR principles provide a framework for data producers to better disseminate and derive value from their data. The consideration of the FAIR principles in policy making in the life sciences was the result of efforts across the wider spectrum of scientific research communities to consolidate good data management practices into a concise set of core values. Whilst these values do not outline the “how”s of data management, but rather the “why”s, they provide a Maslow’s Hierarchy of Needs ^5^ for researchers to start thinking more concretely about their data and their responsibility for its reuse by others, often well beyond the timeframes and immediate purposes originally envisaged when generating it. As such, advancing FAIR requires not just technical, but also cultural, considerations.

For example, as discussed above, the use of defined ontology terms to annotate data and data services helps to render data **finadable and interoperable** - computers can be used to search, aggregate, and integrate data based on the occurrence of compatible terms (e.g. overlapping concepts or synonyms) between datasets. However, life science datasets tend to be disparate, except for a few large, curated, public data repositories (e.g. INSDC data warehouse resources, FlyBase, WormBase, Ensembl Genomes, etc.) The databases and data hosting services that house them may not mandate ontology terms for data description upon submission. Nor may publishers or funders, even those with mature Open Access policies that govern data as well as other research outputs. Furthermore, funding for data resources often comes at the national or research organisation level and historically only limited funding (e.g. the European Union Horizon 2020 calls for e-Infrastructure) has been available to develop global, or integrate international, standards or tools and systems to enhance and exploit them. However, a consensus is growing within the scientific community to make sure data services describe and expose their data using agreed standards ^6^, in spite of the limited funding that may be available ^7^.

From a recent NSF survey, it seems researchers are becoming more mindful of the value of well-described open and FAIR data and their responsibilities to provide it, but they often feel inadequately trained to do so ^8^. The quantity of data produced from scientific experiments is becoming increasingly large and diverse. New approaches, such as BioSchemas ^9^, are attempting to exploit the ubiquity of internet search engines and their metadata harvesting and indexing capacity to expose data through standard search results. Once datasets are discoverable, interoperability then relies on exposure of data through standard Application Programming Interfaces (APIs) of which there are active and notable examples in the life sciences ^10,11^. However, data management tools, especially those used to annotate and describe data assets, tend not to be designed for typical biologists. Rather they are often designed for the small number of professional biocurators, and this may deter other researchers whose participation is essential if curation capacity is to the scale to match the growth of data production. Scientific institutions often lack these biocuration experts who could advise researchers ^12^. Moreover, the variety of research data means it is also important to maximise the pool of software engineers developing biocuration systems. This can be best achieved if code is published using an open-source licence.

### Researchers need help to biocurate

Recording metadata for structured FAIR data remains difficult for researchers, and represents the “last mile” in scientific data publishing ^13^. We believe the solution is to help users to publish FAIR data through user-friendly systems and workflows explicitly designed for that purpose. Alongside these, training and guidance about the ontologies suitable to annotate their data, and repositories suitable to house it, should be provided.

COPO is a platform for researchers to publish their research assets, providing metadata annotation and deposition capability. It allows researchers to describe their datasets according to community standards and broker the submission of such data to appropriate repositories whilst tracking the resulting accessions/identifiers. These identifiers can be used as citations for the richer context of a researcher’s output (i.e. Data production, Images, Source Code etc). As such, COPO directly addresses the Accessible and Interoperable tenets of FAIR ^14^. This has been achieved by developing wizards built into a web-based platform to allow users to describe their data without having to resort to complex spreadsheets, multiple redundant data descriptions when handling multiple research object entities, or multiple submissions to different repositories. These wizards facilitate the use of metadata required by individual repositories, compliance with broader community standards, and are presented to users in an incremental linear fashion so as not to overburden them with large numbers of metadata fields at once. Metadata captured in these wizards is then automatically transformed into the format specified by the repository, removing this task from the user. The design is distinct from that of Dendro ^15^ and CEDAR ^16^, two other platforms with related capabilities. Dendro provides linear web forms. CEDAR uses dynamically created templates and, like COPO, allows arbitrary use of ontologies for data description. However they do not provide rich wizard functionality based on bespoke institutional research data management policies. COPO uses these modular configurable wizard elements based on standard information interchange formats to build complex but easily navigable UIs.

COPO mediates submission to many of the most widely repositories in the life science community (e.g. the EMBL-EBI European Nucleotide Archive for genomics researchers, see Supported Schemas and Repository Types). As it is open source, COPO can be made specific to a community and/or repository used by that community through implementing new wizards that precisely meet their needs. We are working on supporting a larger set of data repositories as COPO matures and as submission routes to these repositories implement documented APIs. We provide two repository Use Cases below.

## Materials and Methods

### Software stack

COPO is primarily a Python application. At the time of writing, COPO runs on a Python 3.5.2 interpreter with Django 2.0.2 as the web framework. Python was chosen as it is a well used and documented language with an active development community. This ensures there are mature libraries for common tasks (such as web development, data loading or ingestion, visualisation, tabular data manipulation, statistical analysis, natural language processing, etc.) as well as cutting edge libraries for modern data science (such as Pandas, scikit-learn, TensorFlow and Numpy/SciPy). Python integrates easily into system-level scripting and is therefore suitable for “glue” projects where many components need to interoperate, such as web service platforms.

The Django web framework was chosen to handle the server side duties of COPO, as it is a tried and tested off-the-shelf solution used by many web services across the international academic, commercial and industrial sectors. A web framework provides developers with many commonly needed functionalities such as URL mapping, page templating, object relational mapping, session handling, user management and security checking. Django is updated regularly, has exceptionally good documentation as well as a large user base.

COPO uses an internal JavaScript Object Notation (JSON) data structure, or *schema*, to store metadata based on modular fragments that can be generated from a range of metadata management processes. JSON is a lightweight, text-based, language-agnostic data interchange format, designed for machines to parse and generate data at scale. JSON is termed a “self-documenting” format, meaning that it is clear enough for a human to understand without reference to any supplementary information. Metadata can be represented in JSON as a collection of name/value pairs. JSON is a common data representation format for many public APIs due to its simplicity and readability. Libraries for manipulating JSON exist for most if not all commonly used programming languages ^17^.

Both Postgres 9.6 and MongoDB 3.4 databases are used for backend storage: Postgres for structured data such as housekeeping tasks related to Django (user accounts, sessions etc); and MongoDB to store the COPO JSON data objects and metadata. MongoDB is a NoSQL database, i.e. the schema does not need to be predefined, facilitating the support of highly heterogeneous metadata without the need for complex schema design. Data files (e.g. raw sequence data, images, PDFs) uploaded to COPO are held on an Isilon storage array - a robust, scale out network-attached storage (NAS) solution for high-volume storage, backup and archiving of unstructured data ^18^. COPO currently benefits from a shared access to over 3.4 petabytes of storage space on this platform, which provides it with the scope to manage the large volume of data brokered through the system. iRODS ^19^ is a framework for abstracting and federating data storage or repositories in a way that promotes data virtualisation, data discovery, data policy automation and secure collaboration across sites. The use of iRODS has been experimented in COPO in order to provide a storage solution that is agnostic of any specific hardware resource architecture.

The user interface (UI) components of COPO have been developed primarily using the Javascript jQuery 2.1.4 library and the Bootstrap 3 framework. Additional javascript UI libraries used include DataTables 1.10.18 for displaying and paginating lists of data, Semantic UI 2.4.0 for UI controls such as buttons and sliders, Handsontable 7.1.0 for displaying spreadsheet type data, and Fuel UX 3.12.0 for creating the COPO wizards.

### Under the Hood

The primary mechanism COPO uses to collect metadata is the *wizard*, which represents the simplest form of an expert system. Wizards consist of a series of interactive stages or form pages which are displayed sequentially, showing users paths through a task based on answers they have given to previous questions. Fields in these wizard stages contain labels, help tips and controls, which guide the user to enter the required metadata. Some of these controls look up bespoke lists of values that are not in a formal ontology but are required by a repository, some (such as calendars) always construct data in a valid format, whilst others perform a lookup call on the Ontology Lookup Service ^20^ and complete a valid ontology reference.

New COPO wizards can easily be configured by defining JSON schemas that comprise the metadata representation for a given repository. Metadata representations between COPO and data repositories are powered by the ISA JSON schema ^21^, allowing information on metadata fields and their rendering to be stored in a central place. For example, fields for a particular type of submission are stored in the same JSON file, with each field having an entry with a unique ID to that field type, specification on type of control to be rendered on a web page to collect the data, whether it’s required, the names it has had in the past, label, help text, etc. (see Figure 2). From this JSON representation, UI controls are automatically generated and rendered to collect such metadata during submission (see Figure 3). This simplifies maintenance and rendering of the UI and also means that creating a submission wizard for a new repository/datatype is as simple as copying and pasting an existing JSON fragment and changing the fields as required. This new configuration will then be rendered without having to write any HTML.

**Figure 2.**
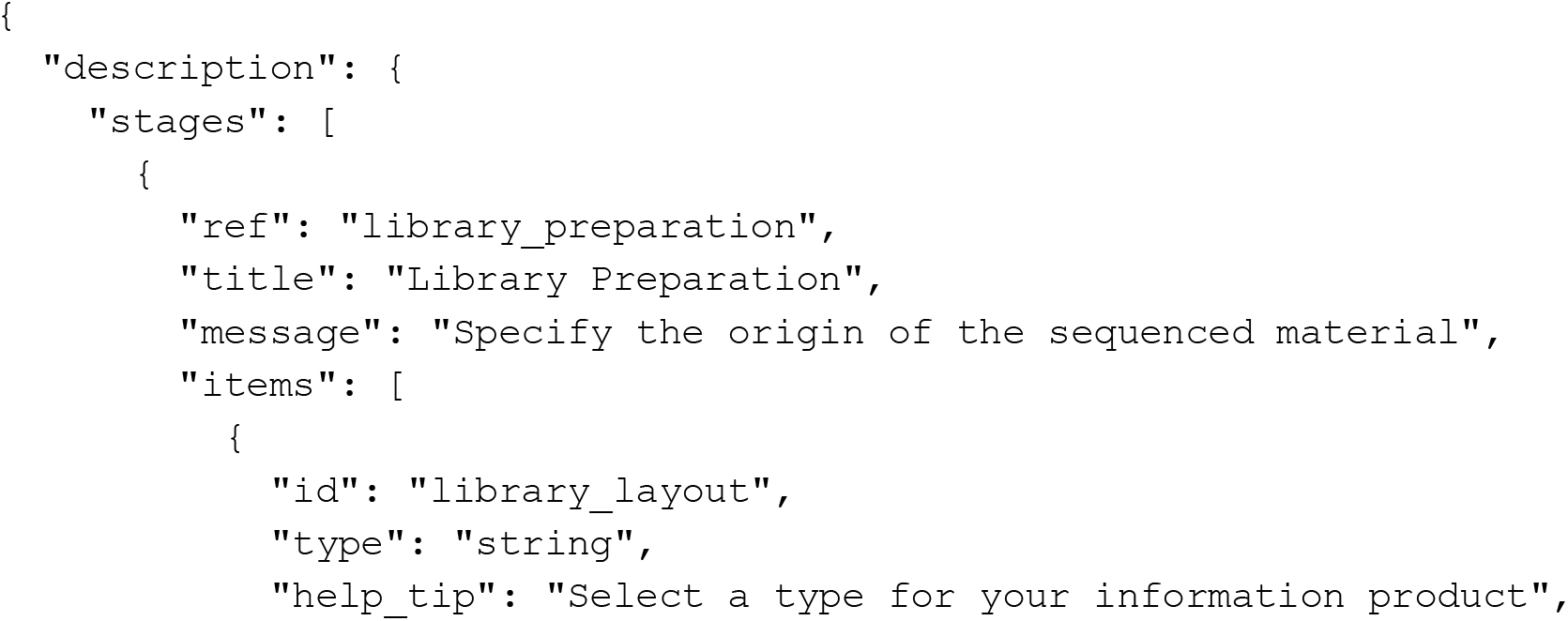
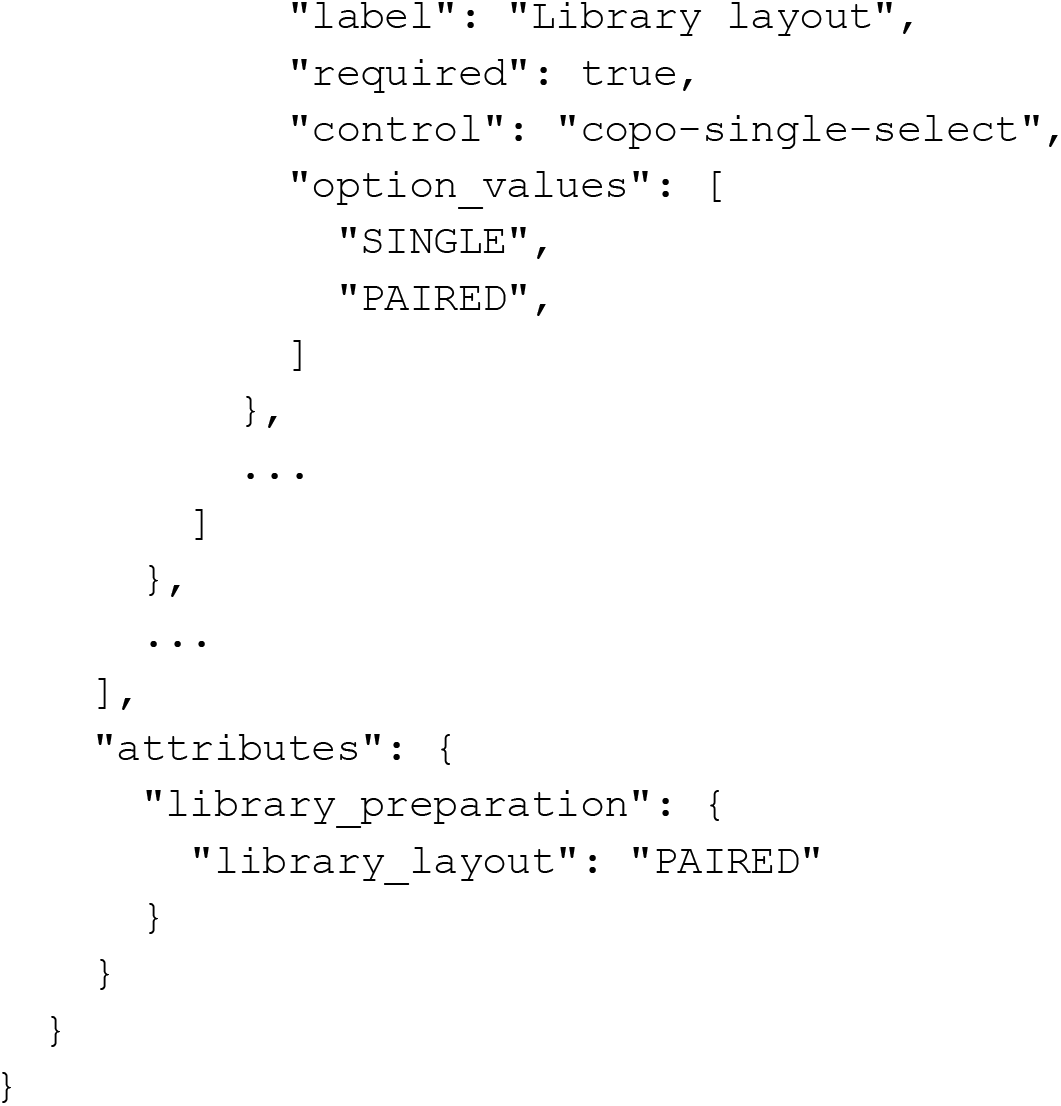
Example COPO schema fragment showing both rendering (under “description”) and data collection (under “attributes”) sections

**Figure 3.**
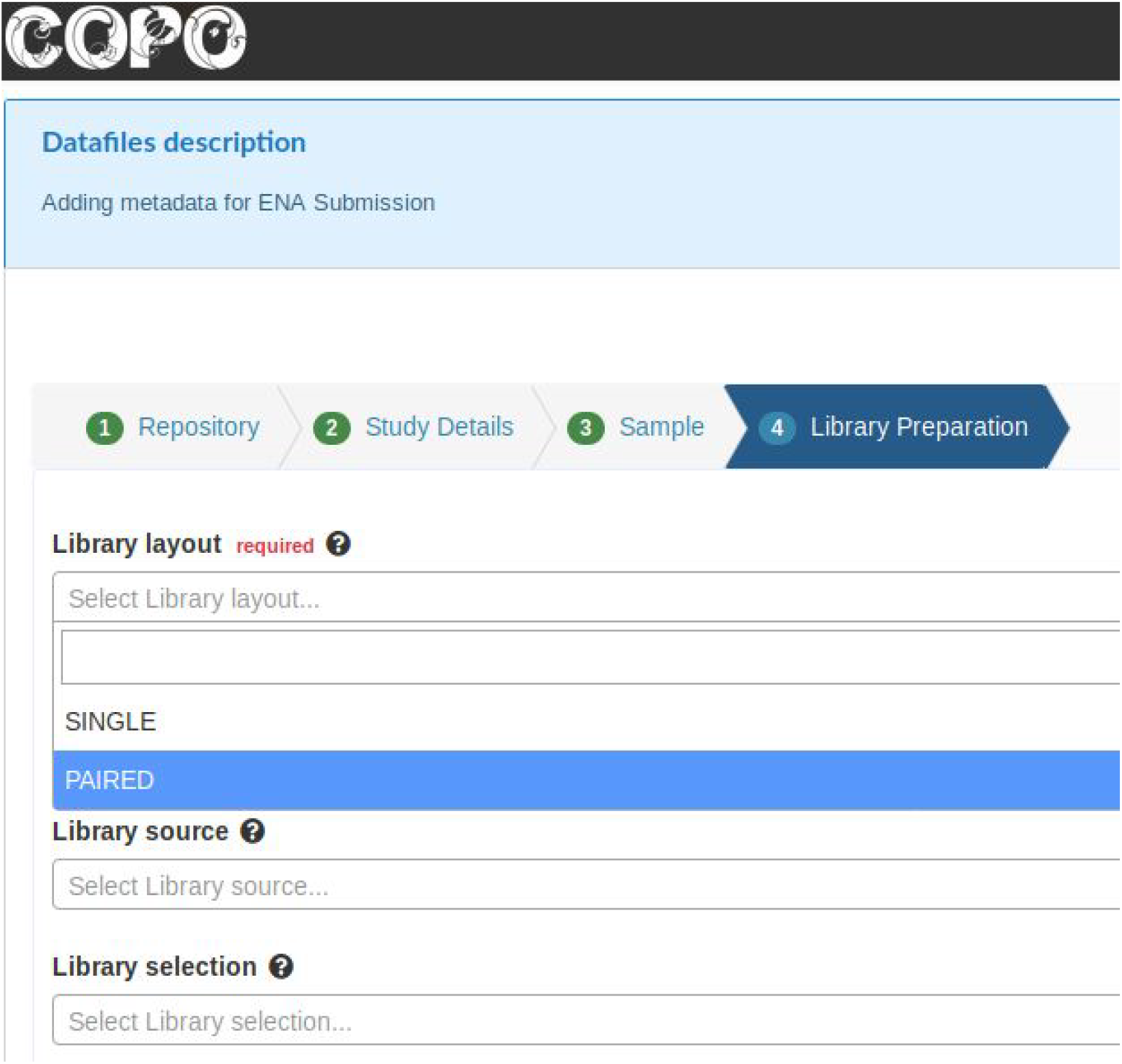
Showing the automatic rendering of the field specified in the schema in Figure 3

### Experimental metadata management

As indicated above, COPO uses the ISA JSON representation to keep experimental metadata for further data sharing and data publication. The ISA framework ^22^ comprises an established data model and a set of open source tools supporting users to provide rich descriptions of their experimental data. ISA stands for Investigation (the project context), Study (a unit of research) and Assay (an analytical measurement). The ISA data model is provided in a variety of serialisations: CSV/TSV tabular format, the Resource Description Framework (RDF) for linked data, and as JSON ^23^. There is a large and international community of ISA adopters and developers associated with the ISAcommons ^22^ that ranges from public repositories to institutional repositories, funded research consortia and data journals.

The ISA model and tools allow the description of experimental details workflows, sample characteristics, technology and measurement types, sample-to-data relationships, and source to sample data processes. The ISA model can also be configured to be compliant with the minimum information requirements for different domains.

### Supported schemas and repository types

#### ‘Omic sequence and feature data

COPO supports the brokering of sequence reads, assemblies, and annotations. We are currently working with repository maintainers to develop wizards for the more esoteric analysis data types, such as optical maps. Data files are uploaded to the EMBL-EBI European Nucleotide Archive servers, and wizards are provided to assist users in fulfilling the requirements of the EBI submission schemas without them needing to create spreadsheets, XML fragments or use the ENA command line tool. The full ENA/SRA metadata fragment is created and the submission happens seamlessly for the user. We describe this in more detail in the Use Case example below.

#### Phenomics data

Collection and deposition of phenotypic experiments is partially supported. The MIAPPE (Minimal Information About Plant Phenotyping Experiments) schema is implemented to collect the required metadata for such experiments. At the time of writing, there is currently no formal repository into which these data can be deposited. However, the EMBL-EBI have recently announced their BioImaging repository ^24^, which is able to accept any image-based phenotyping datasets. Submission will be supported in an upcoming COPO version.

#### Agriculture data

The CG Core metadata schema ^25^ is implemented in COPO as a pilot for agricultural data annotation for the 15 Consultative Group on International Agricultural Research (CGIAR) agricultural research for development centers. A more generalised version of this implementation will soon be more widely available for other entities wanting to describe agricultural data. CG Core is a minimum metadata specification based on the Dublin Core standard to collect descriptive metadata on agronomic, plant breeding, socioeconomic, genomic, and other agricultural information products. Each of the 15 centres store their datasets and publications on centre-specific publicly accessible digital repositories based on one of three repository software implementations, i.e. DataVerse, dSpace or CKAN (see below). The CG Core schema allows consistent description of these data assets for easier discovery and access. We describe this in more detail in the Use Case section.

#### Other scholarly data types

COPO can be used to annotate other information types such as images, presentations, figures, text documents, and PDFs. The Dublin Core metadata specification ^26^ is also employed for these, and submission is supported to institutional instances of Dataverse ^27^, DSpace ^28^ or CKAN ^29^, all of which are free, open-source, off-the-shelf repository applications. COPO also enables submissions to figshare ^30^. Figshare is a centralised general purpose repository whilst CKAN and Dataverse can be installed on local servers (figshare also supports institutional installations of the software but only through a paid model ^31^). There is a publicly accessible instance of Dataverse available for those who cannot host their own ^32^.

### Deployment architecture

The COPO infrastructure is currently housed within the CyVerse UK virtual infrastructure ^33^, supported by the Earlham Institute (EI) National Capability in e-Infrastructure (EI NCG NC3) ^34^. COPO uses the Docker ^35^ containerisation software to house the service deployments, i.e. MongoDB, iRODS, the Django web application and the project website. These Docker containers are managed using Docker Swarm technology to ease orchestration, deployment, and load-balancing challenges (see Figure 4). A default swarm configuration is supplied with COPO, but deployment will need certain properties set by a system administrator, i.e. COPO has a central management node that handles the configuration and setup of the swarm, where system administrators need to supply storage volumes, secret keys, and service definitions for swarm worker nodes. These worker nodes comprise the underlying software stack of databases, web servers and indexers. This modern modular approach to deploying web services makes it possible for third parties to quickly and easily initiate their own versions of COPO.

**Figure 4.**
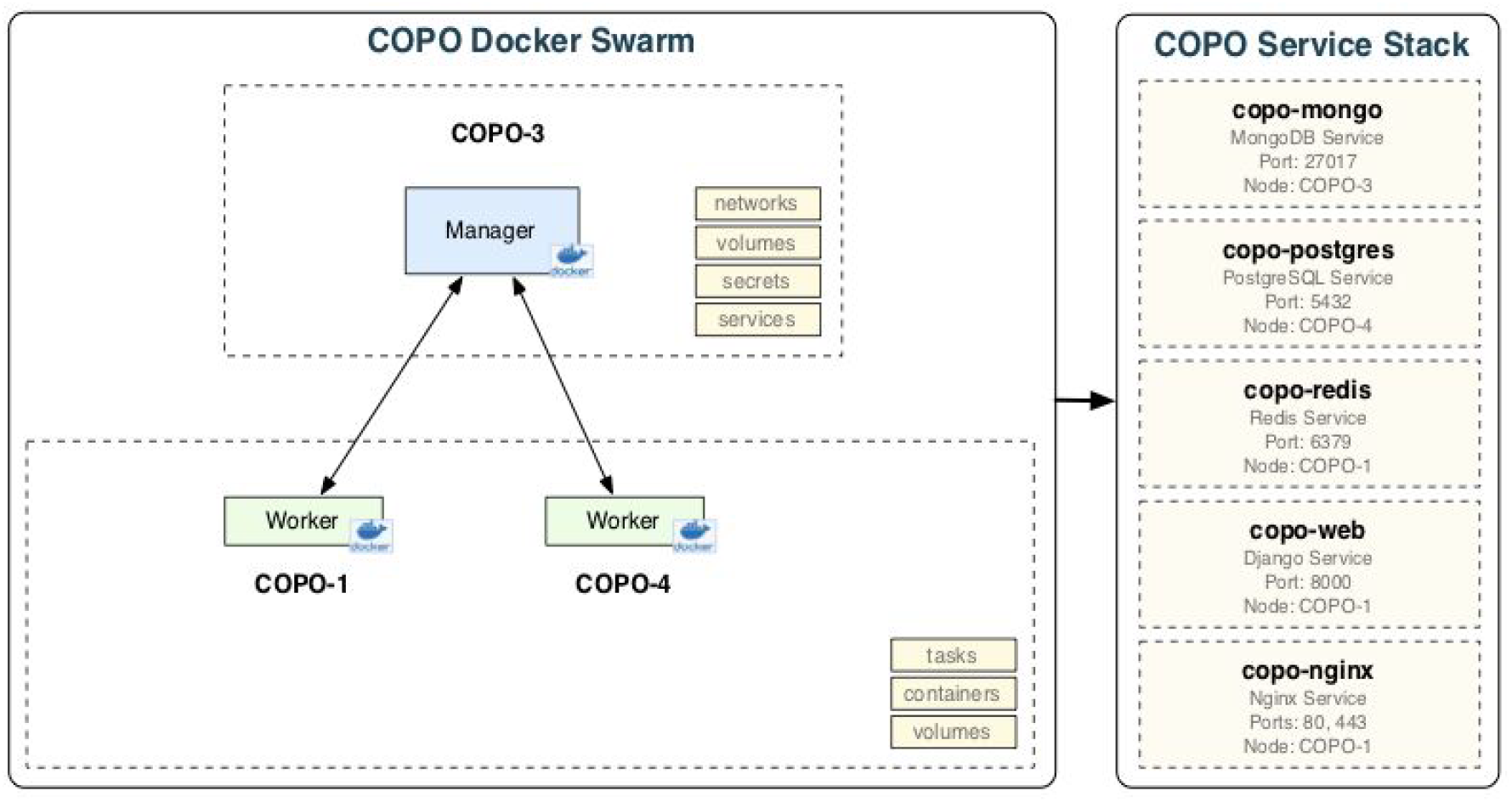
COPO Deployment Architecture using Docker Swarm

### Use Case - Sequence Read Submission to European Nucleotide Archive

A researcher has undertaken a sequencing experiment, and has a number of files containing the filtered and quality-controlled reads representing the nucleic acids that have been sequenced. Once the researcher is ready to submit this read data, they can log into the COPO system using their Open Researcher and Contributor ID (ORCiD). Once logged in, the user can either upload their read data, or describe their samples. UIs have been developed to streamline this process as much as possible. For example, multiple sample objects can be created at once. If the user has existing sample objects in the system, these can be cloned, saving time and effort if a researcher is working on similar organisms, studies, or protocols. Organism names are auto-completed with reference to the NCBI Taxonomy ontology ^36^. Once the required sample objects have been created, the user uploads their sequence data. COPO will compress (gzip) data files if this has not already been done.

The user can then choose a number of files to describe simultaneously in a “bundle”, whereby bundled files are labelled with the same metadata. For example, if there are 50 samples and 50 corresponding files in a bundle, the user only has to go through the description wizard once. Users can also work with unbundled datasets which is useful if the metadata is heterogeneous, e.g. if each data file being submitted is from a different organism or sequenced using different sequencing technologies. The metadata fields which are displayed in the wizard are constructed in line with the schema of the target repository for which the submission is intended. In doing so, the burden of managing metadata revisions, as well as the minimum set of requirements, from different repositories is removed from the researcher and offloaded to COPO. Once the metadata description wizard is complete, the user can then view the metadata for all the files and change any values which are not common in a spreadsheet-like tabular user interface. In the wizard the user also defines which corresponding files each sample object relates to. This is a time-consuming task for researchers as care has to be taken to ensure samples are exactly matched to output files, and COPO aims to reduce errors by making simple sanity checks on sample attribution to data files (number of samples per file, duplicated samples, R1 and R2 files for paired end sequencing experiments, etc). Upon completion, the user is taken to the submission page, where they are shown a progress indicator. During the submission process, the required metadata (in this case ENA submission, study, experiment, run and, as required, analysis XMLs) is produced, the datafiles are physically transferred to the ENA servers, the metadata is validated, and the submission completed (see Figure 5). To our knowledge, COPO is one of the first web-based service implementations of the new EMBL-EBI ENA command line validation client ^37^. The validation process is very resource intensive and the public COPO instance uses the CyVerse UK HPC cloud environment ^33^ to undertake validation.

**Figure 5.**
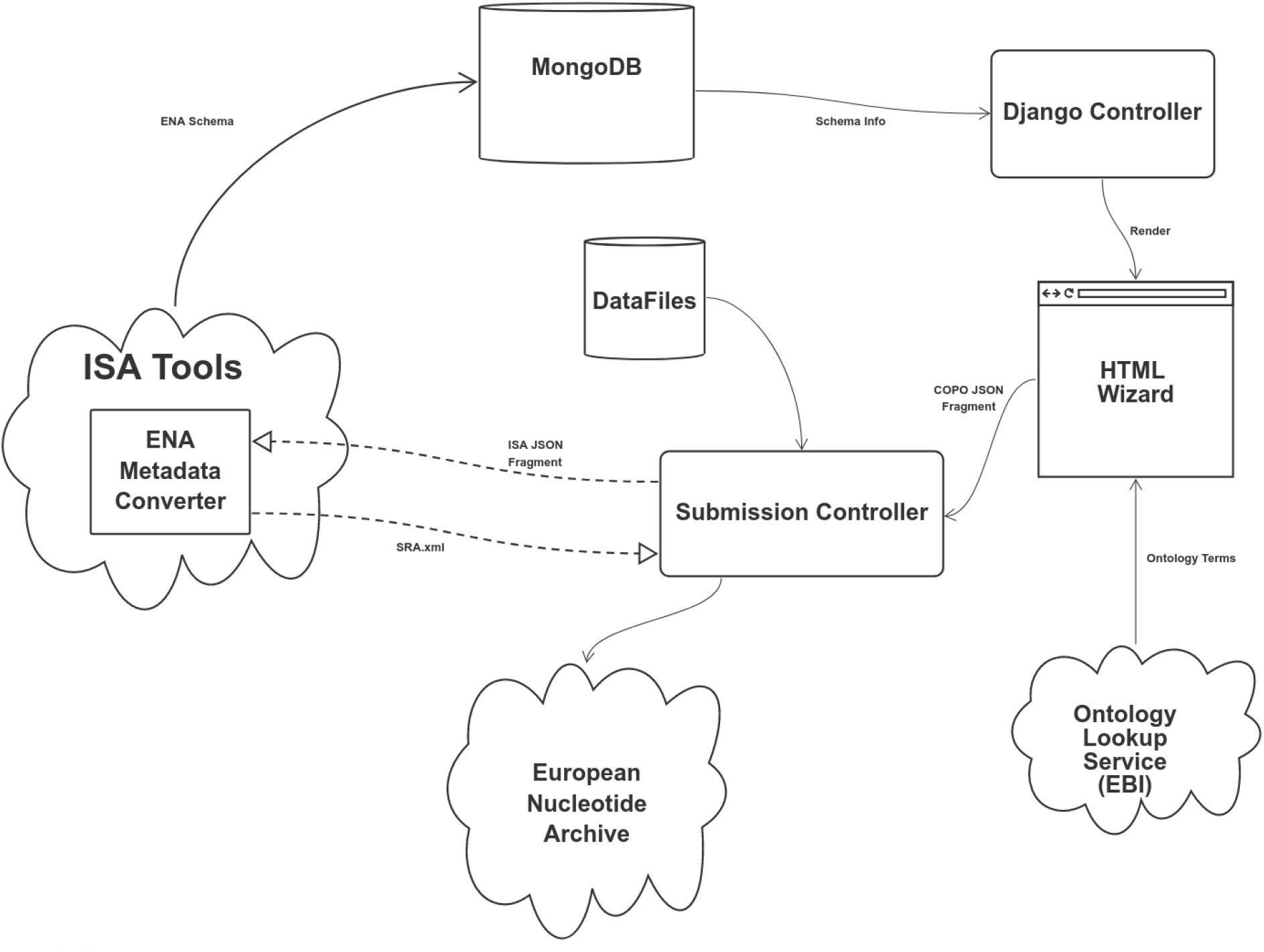
Overview of COPO to ENA Submission Workflow

COPO supports the ENA embargo process whereby researchers can upload data and metadata but not yet make the data public, typically waiting for a publication to be submitted. In any case, ENA accessions for the submission are stored in COPO and are available to the user to publish, and for others to search once public. An example COPO-submitted ENA project can be found under the accession PRJEB30905.

### Use Case - Submission of CGIAR Data to Custom Data Repositories

Digital repositories used by CGIAR centres are hosted locally in some cases, and centralised in others, depending on the repository platform and the centre. Metadata for information resources (typically datasets and publications) are entered by the different data curators and information specialists in each centre. The objective of the CG Core metadata schema is to harmonise descriptions of agricultural resources across the CGIAR centres through a minimum metadata set to maximise discoverability and interoperability. COPO provides a unique entry point to the different systems to guarantee that the metadata are captured in a unified way across the different centre repositories. COPO user interfaces are generated based on the CG Core specification, with the advantage of providing data validation checks while the metadata are entered by the end user. COPO captures the full metadata set of the CG Core, although some data repositories only accept a subset of it. By retaining this metadata alongside links to these repositories, COPO improves the provenance of these records despite the full CG Core metadata not being supported by institutional repositories themselves. This is a by-product of repositories effectively being legacy systems, where either the repository wants to remain general to reduce metadata requirement overload, or highly specific but not able to future-proof any and all possible metadata specifications arising from a community. Managing this extra metadata becomes important for specific internal reporting, where metadata needs to be recorded in a harmonised way, i.e. donor information. Thus in future, as COPO is not intended to be a repository in its own right, we will also submit these additional metadata fragments to a public repository, e.g. Zenodo ^38^, to mint DOIs for the metadata and support data citation (where data is published with a unique identifier, a short publication is written to describe that dataset, and a DOI is minted to allow citation of that dataset to demonstrate use). The full metadata information can then be used by GARDIAN ^39^, the CGIAR metadata harvester and aggregator that allows users to discover datasets and publications using rich queries. The integration of COPO and GARDIAN aims to improve a user’s experience to search across the CGIAR information products (see Figure 6).

**Figure 6.**
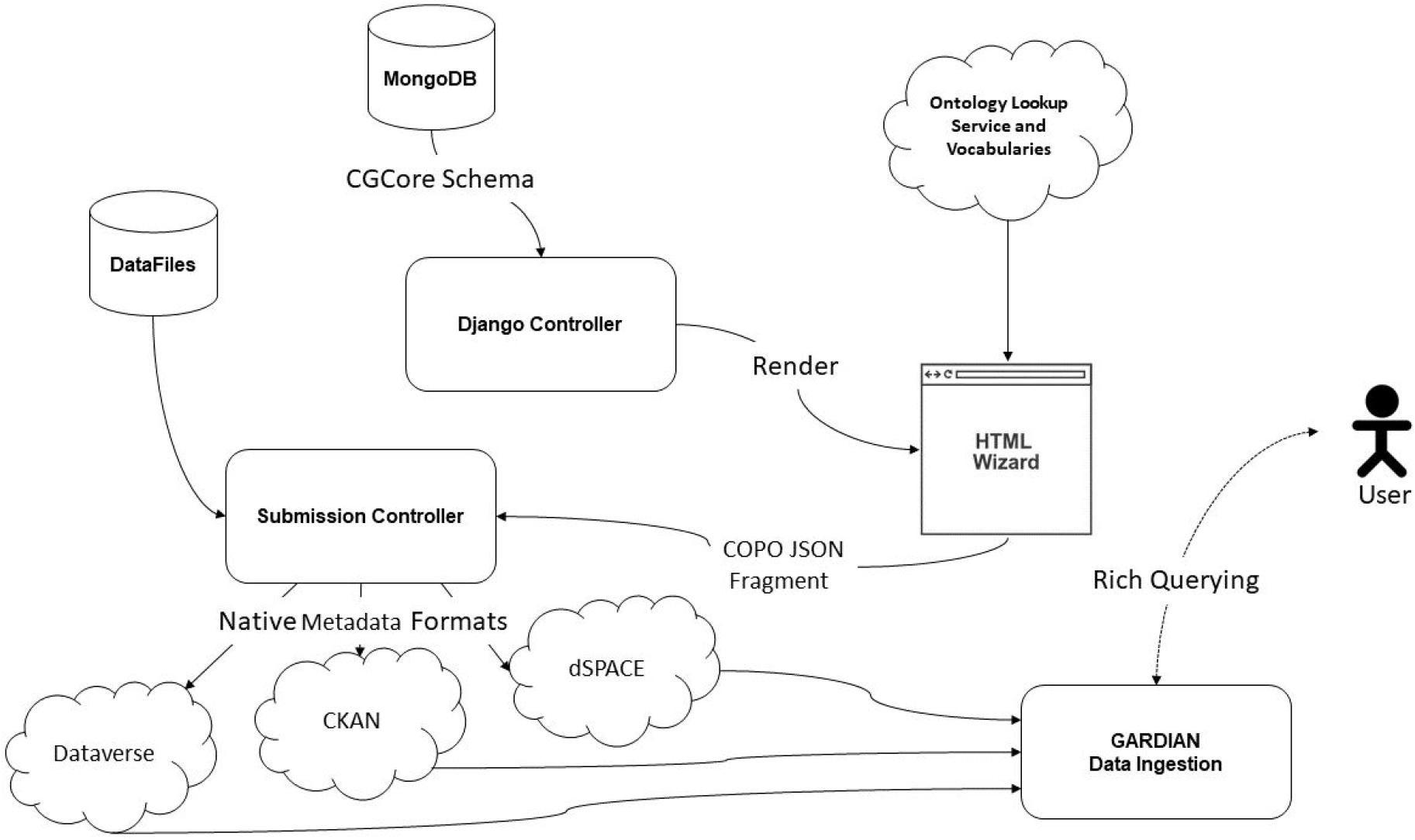
Overview of COPO to CGCore Submission Workflow

## Discussion

The “data deluge” has preoccupied data scientists and knowledge managers for the last decade, has been the subject of popular news articles ^40^, some of which suggest Big Data will see the end of hypothesis-driven science ^41^. The current investment into data generation far outstrips investment in data curation. Many commentaries about the reproducibility and reuse of code, data, and information ^42,43,44,45,46,47^ within publications make sensible points: researchers need to make their digital research outputs available, with requisite associated metadata for reproducibility, and in formats that are open and computationally accessible. However, many if not all of these studies mention in passing, and omit concrete details and processes for, one crucial step: *researchers find it hard and time-consuming to annotate their data with standard vocabularies*. Both technical solutions and coordinated community efforts will be required to manage future datasets and the legacy data that resides in individual group workstations and databases which are no longer actively maintained or curated.

The open source movement in the life sciences is a powerful force, with tools and algorithms developed continuously and released via websites such as Github and Bitbucket. Funders are particularly interested in the impact of algorithms, software tools, and databases to further bioscience. These, including web-based systems, can be submitted as first-class research outputs in many grant reporting platforms, e.g. the United Kingdom Research and Innovation (UKRI) ResearchFish ^42^ system. However, data citation is still a novel concept, and the cost of curation is perceived to be high relative to the cost of producing datasets, particularly raw data generation such as nucleic acid (genome, transcriptome, etc) sequencing. Easing the “FAIRification” of data and enabling the tracking of its impacts can be invaluable for their predestined communities. Further work will enable a rich set of search APIs within COPO to allow users to more accurately track where their data has been used.

The correct ontologies and terms to use to annotate data might not be immediately obvious to researchers, nor where the data should be deposited. FAIRsharing ^48^ works to fill this gap by providing concrete guidance and tools to enable discovery and use of the appropriate terminology, metadata and data standards. It can also help users determine repositories that implement those standards and which journal and funder policies recommend them. However, individual researchers must still search for and assimilate this information. Finding the relevant standards and repositories is only the first step, and institutions may employ dedicated research data managers to help researchers take the next step of annotating data using community standards. Data management employment positions are rare and very specialised to a given domain of knowledge. *Therefore, there remains a professional chasm between researchers who know they need to manage their research data, the tools that are available to allow them to do it, and the people with the epistemological expertise to assist them* ^49^.

Data brokering systems that can support the annotation of data files with rich metadata and submit these easily to data repositories can help greatly to bridge this gap and help researchers realise the practical aspects of sharing FAIR data. Twelve years of experience developing the ISA framework has shown us that standards-driven annotation and data reformatting for brokering purposes needs to be “made invisible” to users. This idea can be brought into line with recent thinking around how to rework the Hierarchy of Needs to something more like a “Hierarchy of Changes” ^50^. COPO is firmly at the “Make it normative” stage, i.e. COPO has been designed with usability in mind, and we have used numerous focus groups to gain feedback on features and the main metadata wizards within the system. We have also presented COPO at events organised by large public data repository managers such as the EMBL-EBI ENA to ensure COPO complements and exploits the submission tools provided by such resources.

Normalising semantic metadata attribution at the researcher and data manager level in COPO means that we now need to consider the “Make it rewarding” and “Make it required” stages. Rewards and incentives for good behaviour help reinforce learning and behavioural change in humans. However, the incentive and reward structures are not yet in place for data (as opposed to journal) publication ^49^. Therefore, to make research data truly FAIR, the scientific community needs to address how sharing and reuse of information outputs is both valued and how it adds value to research in a particular domain. Crop breeding, for example, is an area where standardisation efforts and FAIR data have huge potential to modernise and revolutionise research and applied technological solutions for food security ^51,52^. *These changes are fundamental to allow the application of modern data integration and interrogation approaches and tools to help unlock the value of knowledge held in disparate research groups, institutions, and communities*.

## Conclusions

COPO is a semantic metadata annotation and data brokering platform, initially designed to ease data curation and submission to public repositories for plant scientists. However, as it allows for the flexible selection, constraint, and use of any public or user-specified ontology, COPO is applicable to any domain of knowledge, and has already been customised for agricultural data assets through work with the CGIAR System for international agricultural research for development. As well as a free-to-use central production installation running at the Earlham Institute, COPO is open source under the MIT licence, and therefore freely usable by any institution for their own research data management needs. The tool is packaged in Docker containers for both easy testing and deployment in production. COPO aims to address the “long tail” of data management, where the final stages of preparing data for publication are an often-overlooked resource-intensive part of the publication cycle.

## Acknowledgements

AE, FS and RPD wrote the main text of this manuscript, and all authors contributed to the preparation and writing. The initial COPO project was funded by a BBSRC Biological and Bioinformatics Resources (BBR) grant (BB/L024055/1, BB/L024101/1, BB/L024071/1) and is now funded by the BBSRC Core Strategic Programme grant awarded to Earlham Institute (BBS/E/T/000PR9817). COPO is hosted within the CyVerse UK academic cloud, funded by BBSRC (BB/M018431/1, BB/R000662/1), and run within EI’s National Capability in e-Infrastructure (BBS/E/T/000PR9814). In addition to the BBSRC COPO grant, SAS, PRS, ABG and DJ were funded by BBSRC (BB/L005069/1), Wellcome Trust (212930/Z/18/Z, 208381/A/17/Z), European Union (H2020-EU.3.1, 634107, H2020-EU.1.4.1.3, 654241, H2020-EU.1.4.1.1, 676559), IMI (116060) and NIH (U54 AI117925, 1U24AI117966-01, 1OT3OD025459-01, 1OT3OD025467-01, 1OT3OD025462-01). SAS is also funded by the Oxford e-Research Centre, Department of Engineering Science of the University of Oxford. The computational hardware that underpins this capability is supported by a BBSRC Core Capability Grant to EI, and managed by the Norwich Biosciences Institutes Partnership (NBIP) Computing Infrastructure for Science (CiS) group at the Norwich Research Park.

## Abbreviations

CGIAR: Consultative Group on International Agricultural Research
COPO: Collaborative Open Plant Omics
FAIR: Findable, Accessible, Interoperable and Reusable
INSDC: International Nucleotide Sequence Database Collaboration
ISA: Investigation / Study/ Assay
MIAPPE: Minimum Information About Plant Phenotyping Experiment
JSON: JavaScript Object Notation

## Notes

https://copo-project.org

https://github.com/collaborative-open-plant-omics

## References

1. International Society for Biocuration. Biocuration: Distilling data into knowledge. PLoS Biol. 16, e2002846 (2018).

2. Gruber, T. R. A translation approach to portable ontology specifications. Knowledge Acquisition 5, 199–220 (1993).

3. openrefine. Available at: http://openrefine.org/. (Accessed: 4th September 2019)

4. Wilkinson, M. D. et al. The FAIR Guiding Principles for scientific data management and stewardship. Sci Data 3, 160018 (2016).

5. Maslow, A. A Theory of Human Motivation. (Lulu.com, 1943).

6. Sansone, S.-A. et al. FAIRsharing as a community approach to standards, repositories and policies. Nat. Biotechnol. 37, 358–367 (2019).

7. Reiser L, E. et al. Sustainable funding for biocuration: The Arabidopsis Information Resource (TAIR) as a case study of a subscription-based funding model. - PubMed - NCBI. Available at: https://www.ncbi.nlm.nih.gov/pubmed/26989150. (Accessed: 4th September 2019)

8. Barone, L., Williams, J. & Micklos, D. Unmet needs for analyzing biological big data: A survey of 704 NSF principal investigators. PLoS Comput. Biol. 13, e1005755 (2017).

9. Michel, F. & The Bioschemas Community. Bioschemas & Schema.org: a Lightweight Semantic Layer for Life Sciences Websites. Biodiversity Information Science and Standards 2, (2018).

10. Abbeloos, R. et al. BrAPI - an Application Programming Interface for Plant Breeding Applications. Bioinformatics (2019). doi:10.1093/bioinformatics/btz190

11. SmartAPI | Building a connected network of FAIR APIs. SmartAPI Available at: http://smart-api.info/. (Accessed: 3rd September 2019)

12. Growing demand for data science leaves Britain vulnerable to skills shortages | Royal Society. Available at: https://royalsociety.org/news/2019/05/data-science-skills-shortages/. (Accessed: 9th July 2019)

13. Gabridge, T. Last Mile: Liaison Roles in Curating Science and Engineering Research Data (RLI 265, Aug. 2009).

14. Leonelli, S., Davey, R. P., Arnaud, E., Parry, G. & Bastow, R. Data management and best practice for plant science. Nat Plants 3, 17086 (2017).

15. Silva, J. R. da, da Silva, J. R., Castro, J. A., Ribeiro, C. & Lopes, J. C. Dendro: Collaborative Research Data Management Built on Linked Open Data. Lecture Notes in Computer Science 483–487 (2014). doi:10.1007/978-3-319-11955-7_71

16. Gonçalves, R. S. et al. The CEDAR Workbench: An Ontology-Assisted Environment for Authoring Metadata that Describe Scientific Experiments. Lecture Notes in Computer Science 103–110 (2017). doi:10.1007/978-3-319-68204-4_10

17. JSON. Available at: https://www.json.org/. (Accessed: 20th August 2019)

18. A Closer Look at the Dell EMC Isilon NAS Storage Platform -- Virtualization Review. Virtualization Review Available at: https://virtualizationreview.com/articles/2018/05/15/dell-emc-isilon-nas-storage-platform.aspx. (Accessed: 24th September 2019)

19. iRODS. Available at: https://irods.org. (Accessed: 24th September 2019)

20. Cote, R. G., Jones, P., Martens, L., Apweiler, R. & Hermjakob, H. The Ontology Lookup Service: more data and better tools for controlled vocabulary queries. Nucleic Acids Research 36, W372–W376 (2008).

21. The ISA-JSON format. Available at: https://isa-specs.readthedocs.io/en/latest/isajson.html. (Accessed: 19th August 2019)

22. Sansone, S.-A. et al. Toward interoperable bioscience data. Nat. Genet. 44, 121–126 (2012).

23. González-Beltrán, A., Maguire, E., Sansone, S.-A. & Rocca-Serra, P. linkedISA: semantic representation of ISA-Tab experimental metadata. BMC Bioinformatics 15, 1–15 (2014).

24. BioImage Archive - a new hub for biological images. Available at: https://www.ebi.ac.uk/about/news/press-releases/bioimage-archive-launch. (Accessed: 24th September 2019)

25. AgriculturalSemantics. AgriculturalSemantics/cg-core. GitHub Available at: https://github.com/AgriculturalSemantics/cg-core. (Accessed: 2nd July 2019)

26. DCMI: Dublin Core. Available at: http://dublincore.org/specifications/dublin-core/. (Accessed: 2nd July 2019)

27. The Dataverse Project - Dataverse.org. Available at: https://dataverse.org/. (Accessed: 2nd July 2019)

28. DSpace - A Turnkey Institutional Repository Application. Duraspace.org Available at: https://duraspace.org/dspace/. (Accessed: 3rd September 2019)

29. ckan. ckan Available at: https://ckan.org/. (Accessed: 2nd July 2019)

30. figshare - credit for all your research. Available at: https://figshare.com/. (Accessed: 2nd July 2019)

31. Reed, R. B. figshare for Institutions. J. Med. Libr. Assoc. 104, 376 (2016).

32. Harvard Dataverse. Available at: https://dataverse.harvard.edu/. (Accessed: 20th August 2019)

33. CyVerse UK – CyberInfrastructure for life science. Available at: http://cyverseuk.org/. (Accessed: 3rd September 2019)

34. National Capability in e-Infrastructure. Earlham Institute (2018). Available at: http://www.earlham.ac.uk/national-capability-e-infrastructure. (Accessed: 3rd September 2019)

35. What is a Container? | Docker. Docker Available at: https://www.docker.com/resources/what-container. (Accessed: 24th September 2019)

36. Federhen, S. The NCBI Taxonomy database. Nucleic Acids Research 40, D136–D143 (2012).

37. European Nucleotide Archive. webin-cli. GitHub Available at: https://github.com/enasequence/webin-cli. (Accessed: 24th September 2019)

38. Zenodo - Research. Shared. Available at: https://zenodo.org/. (Accessed: 24th September 2019)

39. GARDIAN. Available at: https://gardian.bigdata.cgiar.org/about.php. (Accessed: 3rd September 2019)

40. Hannay, T. Stop the deluge of science research. the Guardian (2014). Available at: http://www.theguardian.com/higher-education-network/blog/2014/aug/05/why-we-should-publish-less-scientific-research. (Accessed: 3rd September 2019)

41. Mazzocchi, F. Could Big Data be the end of theory in science? Available at: https://www.embopress.org/doi/full/10.15252/embr.201541001. (Accessed: 3rd September 2019)

42. Researchfish: Research Impact Assessment Platform. researchfish Available at: https://www.researchfish.net. (Accessed: 3rd September 2019)

43. Mazzocchi, F. Could Big Data be the end of theory in science? Available at: https://www.embopress.org/doi/full/10.15252/embr.201541001. (Accessed: 3rd September 2019)

44. Six factors affecting reproducibility in life science research and how to handle them. Available at: http://www.nature.com/articles/d42473-019-00004-y. (Accessed: 3rd September 2019)

45. Chen, X. et al. Open is not enough. Nat. Phys. 15, 113–119 (2018).

46. den Beek Jeremy Goecks Rolf Backofen Anton Nekrutenko James Taylor, B. G. J. C. J. K. R. D. N. S. M. Practical Computational Reproducibility in the Life Sciences. Available at: https://www.sciencedirect.com/science/article/pii/S2405471218301406. (Accessed: 3rd September 2019)

47. Fanelli, D. Opinion: Is science really facing a reproducibility crisis, and do we need it to? Proc. Natl. Acad. Sci. U. S. A. 115, 2628–2631 (2018).

48. FAIRsharing. Available at: https://fairsharing.org. (Accessed: 3rd September 2019)

49. Leonelli, S. What Difference Does Quantity Make? On the Epistemology of Big Data in Biology. Big Data Soc 1, (2014).

50. Strategy for Culture Change. Available at: https://cos.io/blog/strategy-culture-change/. (Accessed: 3rd September 2019)

51. Ribaut, J.-M. & Ragot, M. Modernising breeding for orphan crops: tools, methodologies, and beyond. Planta 250, 971–977 (2019).

52. Pommier, C. et al. Applying FAIR Principles to Plant Phenotypic Data Management in GnpIS. Plant Phenomics 2019, 1671403 (2019).

